# Peyer’s patches are a niche for antibiotic-driven expansion of Crohn’s disease–associated adherent-invasive *Escherichia coli*

**DOI:** 10.64898/2026.05.20.726557

**Authors:** Katarina R. Iacobucci, Dominique Tertigas, Aline A. Fiebig, Megan T. Zangara, Michael G. Surette, Brian K. Coombes

## Abstract

Antibiotic exposure is a significant risk factor for Crohn’s disease, yet the tissue-specific consequences of antibiotic-driven dysbiosis remain poorly defined. Adherent-invasive *Escherichia coli* (AIEC), a pathobiont enriched in Crohn’s disease, expands following antibiotic treatment, but whether discrete mucosal niches support this expansion is unknown. Peyer’s patches are specialized lymphoid structures that coordinate mucosal immunity and are frequently associated with early disease lesions, suggesting they may represent a vulnerable site for pathobiont colonization. Here, we show that vancomycin disrupts the Peyer’s patch-associated microbiome, creating a permissive niche that is selectively exploited by AIEC and associated with focal inflammation. Antibiotic treatment markedly increased AIEC burden within Peyer’s patches. AIEC localized within the lymphoid follicle was accompanied by focal tissue pathology and a distinct cytokine signature. In contrast, expansion of resident *E. coli* in the absence of AIEC did not elicit comparable inflammation, indicating that the pathogenic traits of AIEC are required to trigger disease-relevant responses in this niche. Supporting this, genetic disruption of flagellin, long polar fimbriae, or antimicrobial peptide resistance in AIEC attenuated Peyer’s patch colonization or inflammation, revealing separable mechanisms governing niche access and immunopathology. Together, these findings identify Peyer’s patches as a previously unrecognized reservoir for antibiotic-driven AIEC expansion and define a localized host–microbe interaction that links dysbiosis to focal intestinal inflammation. These results provide a mechanistic framework for understanding how antibiotic exposure may precipitate site-specific pathology in Crohn’s disease. Further, these findings highlight that mucosal lymphoid tissues should be considered when evaluating microbiome-targeted therapeutic interventions in Crohn’s disease.

## INTRODUCTION

Crohn’s disease (CD) is a chronic inflammatory disorder of the gastrointestinal tract with rising global incidence and substantial negative impact on patient quality of life (1, 2). Although current therapies can mitigate symptoms, they do not address the underlying drivers of disease, which remain incompletely defined. Increasing evidence points to the gut microbiome as a central contributor to CD pathogenesis, with patients exhibiting reduced microbial diversity and an expansion of mucosal-associated Enterobacteriaceae (3). Among these, adherent-invasive *Escherichia coli* (AIEC) has emerged as a key pathobiont linked to CD (4). *E. coli* is enriched in the mucosa of patients at disease onset (5, 6) and this dysbiosis is exaggerated with antibiotic consumption (6). In addition, tissue-associated AIEC correlates with the severity of ileal disease and was restricted to the inflamed mucosa (7). Together, these observations support a model in which tissue-associated AIEC expansion contributes to both the initiation and progression of intestinal pathology.

Environmental factors that perturb the microbiome can further amplify AIEC-associated disease processes (8). Antibiotic exposure is a major risk factor for CD (9, 10), and is known to disrupt colonization resistance and metabolic functions in the gut, enabling blooms of Enterobacteriaceae (11, 12). Consistent with this, we and others have shown that several different antibiotics, including vancomycin, promote AIEC gut expansion in mouse models, accompanied by increased tissue pathology and sustained mucosal colonization even after treatment cessation (13). These findings raise the possibility that discrete mucosal niches support AIEC persistence and expansion following antibiotic-induced dysbiosis. Peyer’s patches are specialized lymphoid structures in the ileum that coordinate mucosal immune surveillance and are frequently associated with early CD lesions. These tissues represent a unique interface between host immunity and the microbiota, integrating antigen sampling, immune activation, and barrier function. Antibiotic exposure has been shown to disrupt Peyer’s patch immune homeostasis, including depletion of Th17 cells and IgA responses, which are critical for maintaining mucosal defense (14). Although AIEC can access Peyer’s patches via M-cell translocation (15, 16), it remains unclear whether antibiotic-induced perturbations sensitize this compartment to pathobiont colonization and expansion.

Here, we examined whether Peyer’s patches serve as a niche for AIEC expansion during antibiotic treatment. Using a murine model of vancomycin treatment and AIEC infection, we examined microbial community structure, bacterial burden, and host responses within Peyer’s patches. We show that antibiotic exposure restructures the Peyer’s patch-associated microbiome, enabling selective expansion and localization of AIEC within this lymphoid compartment, with attendant focal inflammation. These findings identify Peyer’s patches as a previously unappreciated reservoir for AIEC expansion during antibiotic treatment.

## METHODS

### Bacterial cultures

Wild-type AIEC strain NRG857c was grown overnight in a shaking incubator at 37°C in LB media supplemented with ampicillin (200 µg/mL) and chloramphenicol (34 µg/mL). The Δ*fliC*, Δ*lpf*, and ΔPI-6 AIEC mutant strains were grown in LB under the same conditions with antibiotic selection as outlined in Table 1. The following day, cultures were washed twice in PBS and diluted in PBS to a final concentration of 2x10^9^ cfu/mL.

### Antibiotic-induced AIEC expansion model

Animal experiments were conducted in compliance with the Canadian Council on Animal Care guidelines and followed protocols approved by the Animal Research Ethics Board at McMaster University. All mice were purchased from Charles River and were housed in the Central Animal Facility at McMaster University in a level 2 Biosafety room. Six-to-eight-week-old female C57BL/6 N mice were pre-treated with 20 mg of streptomycin via oral gavage. The following day, mice were colonized with 100 µL AIEC NRG857c (2 x 10^9^ cfu/mL) via oral gavage. Uninfected control groups were given 100 µL PBS via gavage. After a five-day colonization period, mice were treated with vancomycin (50 mg/kg) via oral gavage daily for five days. Untreated groups were given a sham treatment of sterile water (5 µL/g body weight) for five days.

### Ileal Peyer’s patch collection and sample processing

At the experiment endpoint, mice were sacrificed via cervical dislocation, and the ileum and Peyer’s patches were harvested. We considered the ileum as the most distal eight centimeters of the small intestine. The luminal contents within the ileum were gently removed with sterile forceps. The ilea were then flushed with approximately 8 mL of PBS to remove remaining contents. Peyer’s patches were extracted from the tissue using curved scissors. All samples were then transferred to pre-weighed Eppendorf tubes. A sterile metal bead was added to ileal tissue and Peyer’s patch samples to aid with tissue homogenization. Luminal content and Peyer’s patch samples were homogenized in PBS for five minutes, while bulk ileal tissue samples were homogenized for 10 minutes.

### AIEC quantification within Peyer’s patches

Tissue homogenates were serially diluted and plated on LB agar supplemented with the appropriate antibiotic selection described above. Plates were incubated overnight at 37°C. The following day, colonies were counted and cfu values were normalized to the weight of the tissue sample.

### Quantification and identification of commensal bacteria within Peyer’s patches

To quantify commensal microbes in Peyer’s patches, tissue homogenates were plated on Brain Heart Infusion (BHI) agar supplemented with hemin (10 mg/L), L-cysteine (0.5 g/L), and vitamin K (1 mg/L). All homogenates were plated in duplicate and grown both aerobically and anaerobically. The anaerobic growth condition (10% CO_2_, 10% H_2_, 85% N_2_) was achieved using the Anoxomat system. BHI plates for both growth conditions were then incubated overnight at 37°C. Colonies were counted and cfu values were normalized to the weight of the sample.

### DNA extraction and 16S rRNA gene sequencing of Peyer’s patches

Peyer’s patches were harvested from mice as previously described and placed in a sterile tube with 1.4 mm ceramic spheres, 0.1 mm silica spheres and one 4 mm glass sphere [mpbio], along with 100 μl thiocyanate guanidinium-EDTA-N-lauroylsarkosine buffer, and 800 μl of 200 mM sodium phosphate monobasic [pH 8] made with UltraPure distilled water [Invitrogen]. Genomic DNA was extracted from Peyer’s patches using a previously described DNA extraction method with minor modifications (17). Here, the sample supernatant was processed using the KingFisher Apex 96-Deep Well Magnetic Particle Processor from Thermo Scientific with the DNA Multi-Sample kit (Life Technologies #4413022). A modified elution volume of 50 µL was used to concentrate the DNA.

A two-step nested polymerase chain reaction (PCR) was used to amplify the variable regions 3 and 4 of the 16S rRNA gene. In the first reaction, 100 ng of purified DNA was combined with 10 pmoles of 8F (AGAGTTTGATCCTGGCTCAG) and 926R (CCGTCAATTCCTTTRAGTTT) primers, 1U of Taq polymerase, 1x buffer, 1.5 mM MgCl2, 0.4 mg/mL bovine serum albumin (BSA), and 0.2 mM deoxynucleoside triphosphates (dNTPs). The reaction was carried out at 94°C for 5 min, then 15 cycles of 94°C for 30s, 56°C for 30s, and 72°C for 60s, followed by a final extension of 72°C for 10 min. In the second reaction, 3 uL of the first reaction was combined with 5 pmoles of 341F (CCTACGGGNGGCWGCAG) and 806R (GGACTACNVGGGTWTCTAAT) Illumina adapted primers(18), 1U of Taq polymerase, 1x buffer, 1.5 mM MgCl2, 0.4 mg/mL BSA, 0.2 mM dNTPs. The reaction was carried out at 94°C for 5 min, then 5 cycles of 94°C for 30s, 47°C for 30s, and 72°C for 30s, followed by 30 cycles of 94°C for 30s, 50°C for 30s, and 72°C for 30s, with a final extension of 72°C for 10 min. PCR products were visualized on a 1.5% agarose gel and amplicons were normalized based on band intensity before sequencing on the Illumina MiSeq system at McMaster Genomics Facility. Raw reads were processed as previously described in DADA2 (17, 19), and the taxonomy of each read was assigned using the SILVA database v138.2 (20).

Microbiome analyses were performed using R v4.4.2. Amplicon sequence variants (ASVs) unclassified at the kingdom or phylum level or ASVs classified as Eukaryota, Archaea, Mitochondria, or Chloroplast were removed. Due to the low microbial load, the prevalence method in decontam v1.26.0 (21) was used to identify contaminants based on a threshold of 0.5 to identify ASVs more prevalent in the PBS control samples compared to the Peyer’s patches samples. Three additional ASVs were removed with a threshold of 0.56 that were present in PBS control samples. Samples containing fewer than 1000 reads were excluded from the analysis. Alpha and beta diversity analyses were performed using Phyloseq v1.50.0 (22) and Microbiome v1.28.0 packages (23) . Beta diversity metrics, including Bray-Curtis dissimilarity and Aitchison distance, were visualized using Principal Coordinates Analysis (PCoA) and Redundancy Analysis (RDA), respectively. The adonis2 function in the vegan package v2.6-10 (24) was used to perform a permutational multivariate analysis of variance (PERMANOVA). Alpha diversity was measured using the Shannon diversity index on data rarefied to the minimum library size, and the significance was determined using a Wilcoxon rank sum test (stats v4.4.2 package). The Wilcoxon rank sum test was also used to assess changes in *Escherichia coli* relative abundance. Results were visualized using tidyverse v2.0.0 (25) and ggpubr v0.6.0 (26) .

### AIEC Immunohistochemistry

Ileal samples harvested at endpoint were flushed of luminal contents, arranged as Swiss-rolls and fixed in 10% neutral buffered formalin for 48 hours. Samples were transferred to 70% EtOH and submitted to the McMaster Immunology Research Centre histology core for embedding and sectioning. Swiss rolls were paraffin-embedded, sliced into 5 µm sections, and mounted onto positively charged slides. Ileal tissues were stained using a standard immunofluorescence protocol. Briefly, slides were de-paraffinized in an alcohol series and placed in antigen retrieval buffer (10 mM sodium citrate, Tween-20, pH 6). A mouse-on-mouse blocking kit was used to prevent endogenous reactivity of the anti-E-cadherin antibody in subsequent steps. Slides were then stained with mouse anti-E-cadherin (1:200; ECM Biosciences CM1681) and rabbit anti-O83 (1:200; SSI Diagnostica, #85077) antibodies overnight at 4°C. The following day, slides were stained with DAPI, anti-mouse Alexa fluor 488 (1:2000), and anti-rabbit Alexa fluor 568 (1:200) secondary antibodies. Coverslips were then mounted and sealed onto slides. Representative Peyer’s patches were imaged using a Zeiss inverted confocal microscope at 20X magnification with Airyscan.

### Histopathology analysis of focal inflammation around Peyer’s patches

At the experiment endpoint, mice were sacrificed via cervical dislocation. The ilea were arranged into Swiss rolls, fixed, sectioned, and embedded as previously described. Sections were stained with hematoxylin and eosin. Slides were then imaged at 10X magnification with a Zeiss slide scanner. Peyer’s patches were located within 10X images of each H&E-stained ileum. The first 5 villi and crypts adjacent to each Peyer’s patch were scored using the histopathology scoring criteria shown in Table 2. Tissue segments on both sides of the Peyer’s patch were examined where possible.

### Cytokine and chemokine profiling in Peyer’s patches

Ileal Peyer’s patches were harvested from mice and homogenized as previously described. Homogenates were centrifuged at 4,000 rpm for 15 minutes at 4°C to pellet particulate matter and debris. The supernatant was collected to quantify total protein levels in each sample using a BCA assay, normalized across samples, aliquoted, and sent for cytokine and chemokine profiling by Eve Technologies (Alberta, Canada). Thirty-six markers were measured in the samples using the Eve Technologies#39; Mouse Cytokine/Chemokine 36-Plex Discovery Assay® Array (MD36) as per the manufacturer’s instructions for use (MILLIPLEX® Mouse Cytokine/Chemokine Magnetic Bead Panel Cat. #MCYT1-190K, MilliporeSigma, Burlington, Massachusetts, USA). Assay sensitivities of these markers range from 0.13– 43.62 pg/mL. Samples are represented using a mean z-score computed using R Studio (Version 2025.09.0+387 (2025.09.0+387)). Outliers were removed following calculation of the interquartile range (IQR).

### Statistical Analysis

Statistical analysis was performed using the GraphPad Prism software (version 10.4.1). Outliers were removed with a q-test where appropriate. Statistical tests were performed as indicated in figure captions. P values less than 0.05 were considered significant.

## RESULTS

### Vancomycin creates a niche for endogenous *E. coli* in Peyer’s patches

Antibiotic treatment has been linked to disruptions in immune activation and function within small intestinal lymphoid compartments such as Peyer’s patches, which play a key role in sensing and responding to microbes. However, it remains unclear whether these changes are accompanied by alterations in the Peyer’s patch-associated microbiome. To test this, we treated naïve specific pathogen-free mice with vancomycin and examined the bacterial burden and endogenous community composition in ileal Peyer’s patches with or without antibiotic treatment. Vancomycin-treated mice had a significant increase in total bacterial load within Peyer’s patches **(Fig 1a)**, suggesting that this compartment becomes more permissive to colonization by endogenous microbes following antibiotic exposure. To determine whether this enrichment reflected broader changes in microbial community structure, we profiled the Peyer’s patch-associated microbiome using 16s rRNA sequencing. The bacterial composition of Peyer’s patches from vancomycin-treated mice was markedly different from non-antibiotic controls (**Fig. 1b)**. In control mice, the Peyer’s patch-associated microbiome was dominated by segmented filamentous bacteria (SFB; *Candidatus arthromitus*), whereas vancomycin treatment led to a contraction of SFB and dominance of *Ligilactobacillus*. Although trending towards reduced diversity following vancomycin treatment, the Shannon diversity index was not statistically different between the Peyer’s patch-associated microbiome of the groups **(Fig. 1c)**. Microbial profiles were significantly different between water and vancomycin-treated mice based on Bray-Curtis dissimilarity (**Fig. 1d**) but compositional differences were not significant by Aitchison distance (**Fig. 1e)**. Because antibiotic-induced dysbiosis can foster the expansion of disease-associated Enterobacteriaceae, including *Escherichia coli* (13, 27, 28) we wanted to understand how vancomycin treatment altered the abundance of endogenous *E. coli*, which is otherwise in very low abundance in the uncontrived microbiome. Vancomycin treatment indeed increased the relative abundance of Peyer’s patch-associated *E. coli* ∼6-fold, however, this did not reach statistical significance (**Fig. 1f**). Together, these findings indicate that vancomycin restructures the Peyer’s patch-associated microbiome and suggested that this altered niche may be permissive to expansion of pathogenic *E. coli* strains such as AIEC.

**Figure 1.**
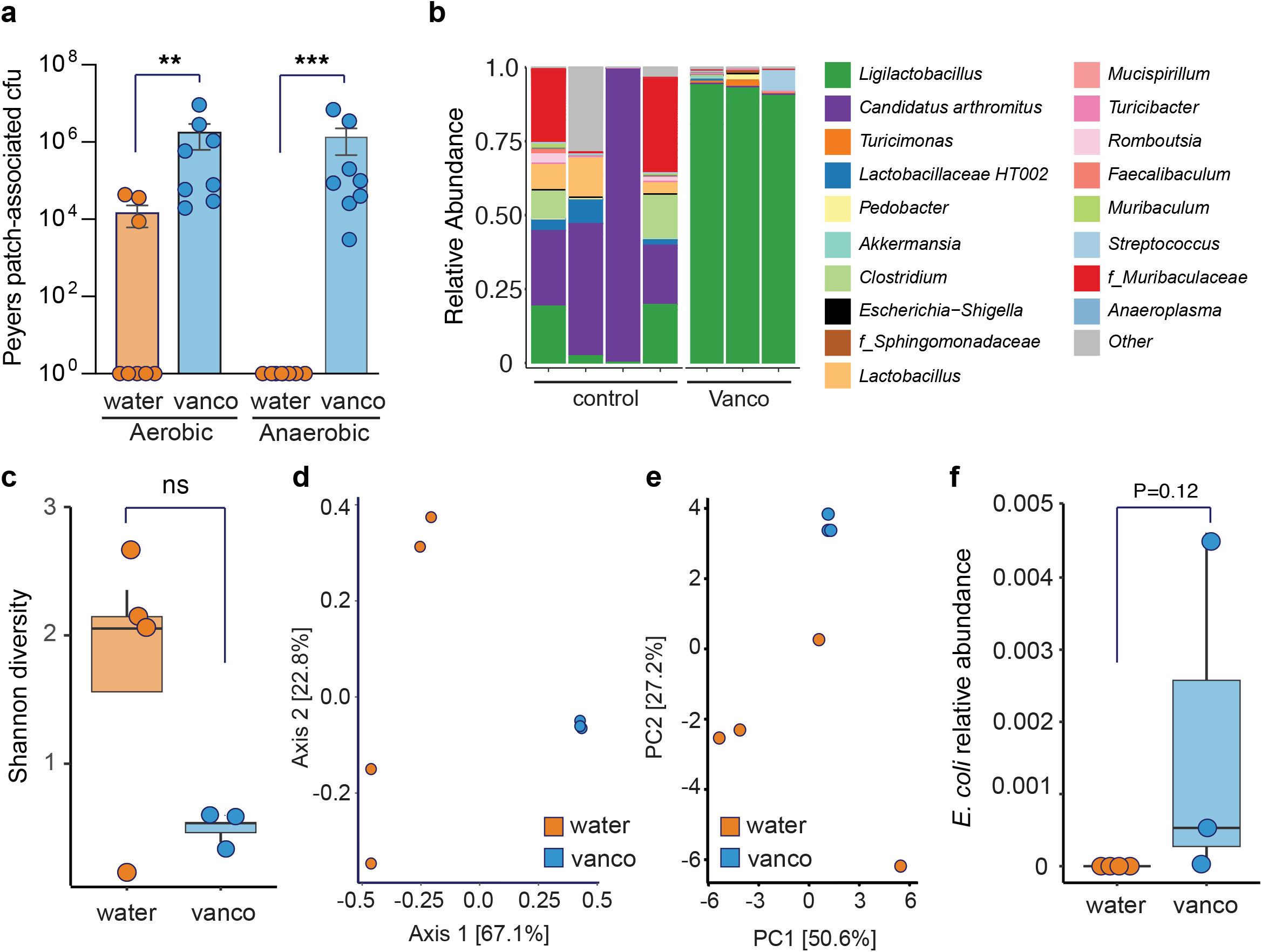
Vancomycin restructures the Peyer’s patch-associated microbiome and promotes endogenous *E. coli* expansion. **(a)** Peyer’s patch-associated CFU from control mice (water) and mice treated with vancomycin for five days under aerobic and anaerobic growth conditions. Bars represent the mean ± SEM with individual mice shown. **p < 0.01, ***p < 0.001 (two-tailed Mann–Whitney test). **(b)** Relative abundance of bacterial taxa at the genus level within the Peyer’s patch-associated microbiome. **(c)** Shannon diversity index of Peyer’s patch-associated microbial communities from water- and vancomycin-treated mice (Wilcoxon rank-sum exact test, p = 0.4). **(d)** Principal coordinates analysis (PCoA) of amplicon sequence variants (ASVs) based on Bray–Curtis dissimilarity (PERMANOVA: R^2^ = 0.64569, p = 0.031). **(e)** Redundancy analysis (RDA) based on Aitchison distance (PERMANOVA: R^2^ = 0.24796, p = 0.216). **(f)** Relative abundance of endogenous *E. coli* within Peyer’s patches (Wilcoxon rank-sum test with continuity correction, p = 0.1227). Each point represents an individual mouse. Bars represent the mean ± SD.

### Vancomycin promotes AIEC expansion in Peyer’s patches

Having established that vancomycin treatment disrupts the endogenous microbiome in Peyer’s patches in naïve mice, including increased abundance of *E. coli*, we next asked whether AIEC would colonize this niche following antibiotic treatment (**Fig. 2a**). Mice colonized with AIEC before vancomycin treatment showed typical levels of luminal and tissue-associated AIEC in the ileum and was undetectable in Peyer’s patches from mice not exposed to vancomycin (**Fig. 2b-d**). In contrast, in mice receiving vancomycin treatment, there was a significant increase in AIEC burden in the ileal lumen, bulk ileal tissue, and a marked increase in Peyer’s patches (**Fig. 2b-d**). AIEC was not detected in Peyer’s patches of every mouse by culture, likely reflecting the low biomass and variable size of these tissues, as some mice possessed only a single or very small Peyer’s patch. To address this limitation, we performed 16s rRNA sequencing to detect Peyer’s patch-associated AIEC. Consistent with the cfu data, the relative abundance of *E. coli* was significantly higher in Peyer’s patches from AIEC-colonized, vancomycin-treated mice compared to controls (**Fig. 2f**). Analysis of the microbiome composition following vancomycin-induced AIEC expansion revealed patterns similar to those observed in naïve mice, including depletion of SFB (**Fig. 2e**). Again, although there was no difference in Shannon diversity between treatment groups (**Fig. 2g**), microbial profiles clustered by treatment group (**Fig. 2h-i**). Together, these findings show that vancomycin-induced dysbiosis is associated with robust expansion of AIEC within Peyer’s patches.

**Figure 2.**
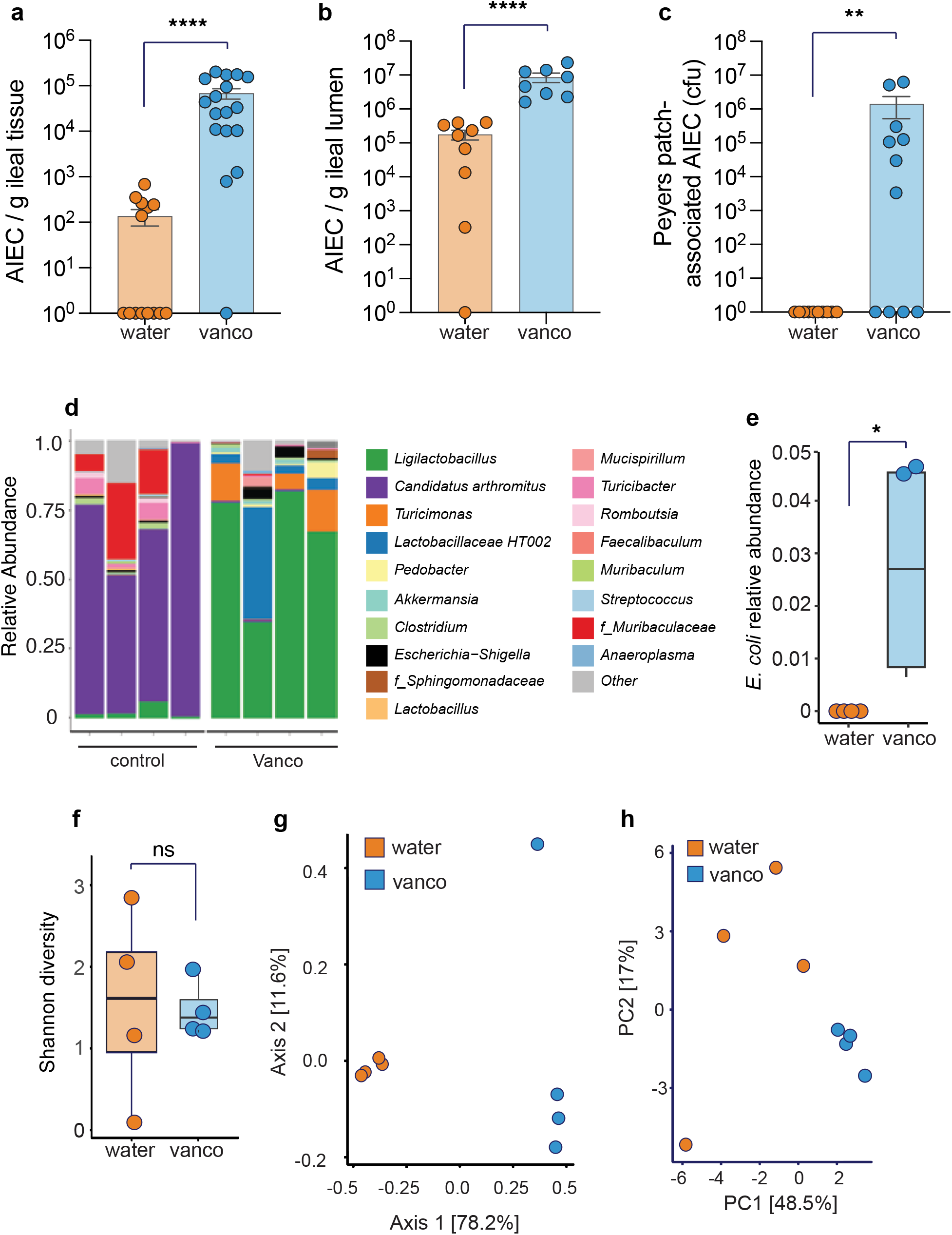
Vancomycin-induced dysbiosis promotes AIEC expansion in Peyer’s patches. **(a)** AIEC burden in ileal tissue, **(b)** ileal lumen, and **(c)** Peyer’s patches from mice colonized with AIEC and treated with either water or vancomycin (50 mg/kg) daily for five days. Each point represents an individual mouse. Bars represent the mean ± SEM. **p < 0.01, ****p < 0.0001 (two-tailed Mann–Whitney test). **(d)** Relative abundance of bacterial taxa at the genus level within the Peyer’s patch-associated microbiome of AIEC-colonized mice treated with water or vancomycin. **(e)** Relative abundance of *E. coli* within Peyer’s patches (Wilcoxon rank-sum test with continuity correction, p = 0.0211). **(f)** Shannon diversity index of Peyer’s patch-associated microbial communities from water- and vancomycin-treated mice (Wilcoxon rank-sum exact test, p = 0.4). **(g)** Principal coordinates analysis (PCoA) of amplicon sequence variants (ASVs) based on Bray–Curtis dissimilarity (PERMANOVA: R^2^ = 0.77865, p = 0.031). Redundancy analysis (RDA) based on Aitchison distance (PERMANOVA: R^2^ = 0.36731, p = 0.03). Each point represents an individual mouse. Bars represent the mean ± SD.

### AIEC localizes within the Peyer’s patch follicular compartment

AIEC are known to adopt multiple lifestyles within the mucosa, including extracellular, intracellular, and in biofilms (4, 29, 30) . We performed immunofluorescence microscopy to visualize AIEC within Peyer’s patches and determine their predominant lifestyle(s) during vancomycin treatment. Consistent with our cfu data, AIEC were not detected in mice from the water-treated control groups (**Fig. 3a**), however AIEC were highly abundant in the Peyer’s patches of vancomycin-treated mice (**Fig. 3b**). Most of the AIEC had breached the follicle-associated epithelium and were found within the lymphoid follicle itself. These infiltrated AIEC existed as single bacteria as well as multicellular foci near the germinal centre of the Peyer’s patch (**Fig. 3d**). We did not observe structures consistent with AIEC biofilms within Peyer’s patches. These observations reinforce that vancomycin promotes AIEC colonization of Peyer’s patches, and that these AIEC primarily reside within the follicular zone of this niche.

**Figure 3.**
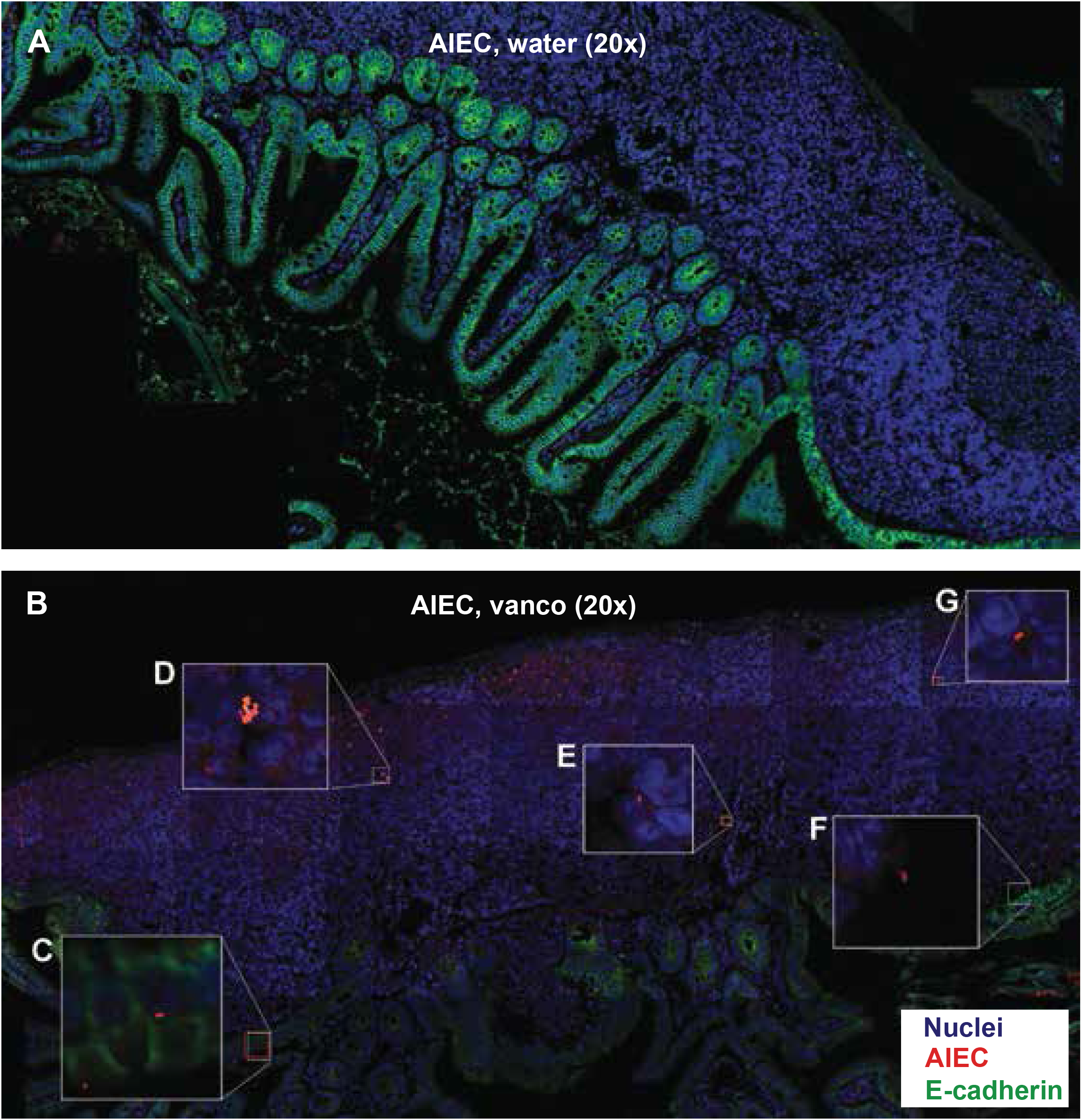
AIEC localizes within ileal Peyer’s patches following vancomycin treatment. Representative immunofluorescence images showing AIEC localization within Peyer’s patches from AIEC-colonized mice treated with either water **(a)** or vancomycin **(b)**. AIEC burden was markedly increased in Peyer’s patches following vancomycin treatment. Insets highlight representative examples of AIEC localization within the follicle-associated epithelium (c), multicellular AIEC clusters within Peyer’s patches (d), and individual AIEC associated with the lymphoid follicle (e–g). AIEC are shown in red, nuclei in blue, and intestinal epithelium (E-cadherin) in green. Overview images were acquired at 20× magnification using AiryScan imaging to improve resolution. Insets represent magnified regions from the corresponding overview image.

### Vancomycin-induced AIEC expansion is associated with focal inflammation around Peyer’s patches

Given that vancomycin promotes AIEC expansion within ileal Peyer’s patches, we next examined the impact of this expansion on the local inflammatory landscape. Ileal tissues were collected from naïve and AIEC-infected mice treated with either water or vancomycin, and histopathological analysis was performed on Peyer’s patch-associated regions. AIEC-expanded mice displayed significantly higher levels of tissue pathology surrounding Peyer’s patches compared to all three control groups (**Fig. 4a– e**). Although vancomycin-treated naïve mice exhibited increased *E. coli* abundance in our microbiome analysis, these mice showed significantly lower pathology scores than AIEC-expanded mice. These findings indicate that AIEC expansion, rather than the presence of resident *E. coli*, is associated with focal inflammation in this compartment during vancomycin treatment. To further assess whether these differences were reflected at the molecular level, we profiled 36 cytokines and chemokines within Peyer’s patches using a multiplex assay (**Fig. 4f**). Vancomycin treatment alone did not significantly alter any of the analytes measured in naïve mice. AIEC infection in the absence of antibiotics resulted in a reduction in IL-2, but did not significantly affect other cytokines or chemokines. In contrast, AIEC-expanded mice displayed the most pronounced changes in cytokine/chemokine profiles. In these mice, we observed a significant decrease in the Th17 cytokine IL-17A, along with reductions in IFN-α and IL-2. Together, these data show that vancomycin-associated AIEC expansion is linked to focal inflammation and distinct alterations in cytokine profiles within Peyer’s patches.

**Figure 4.**
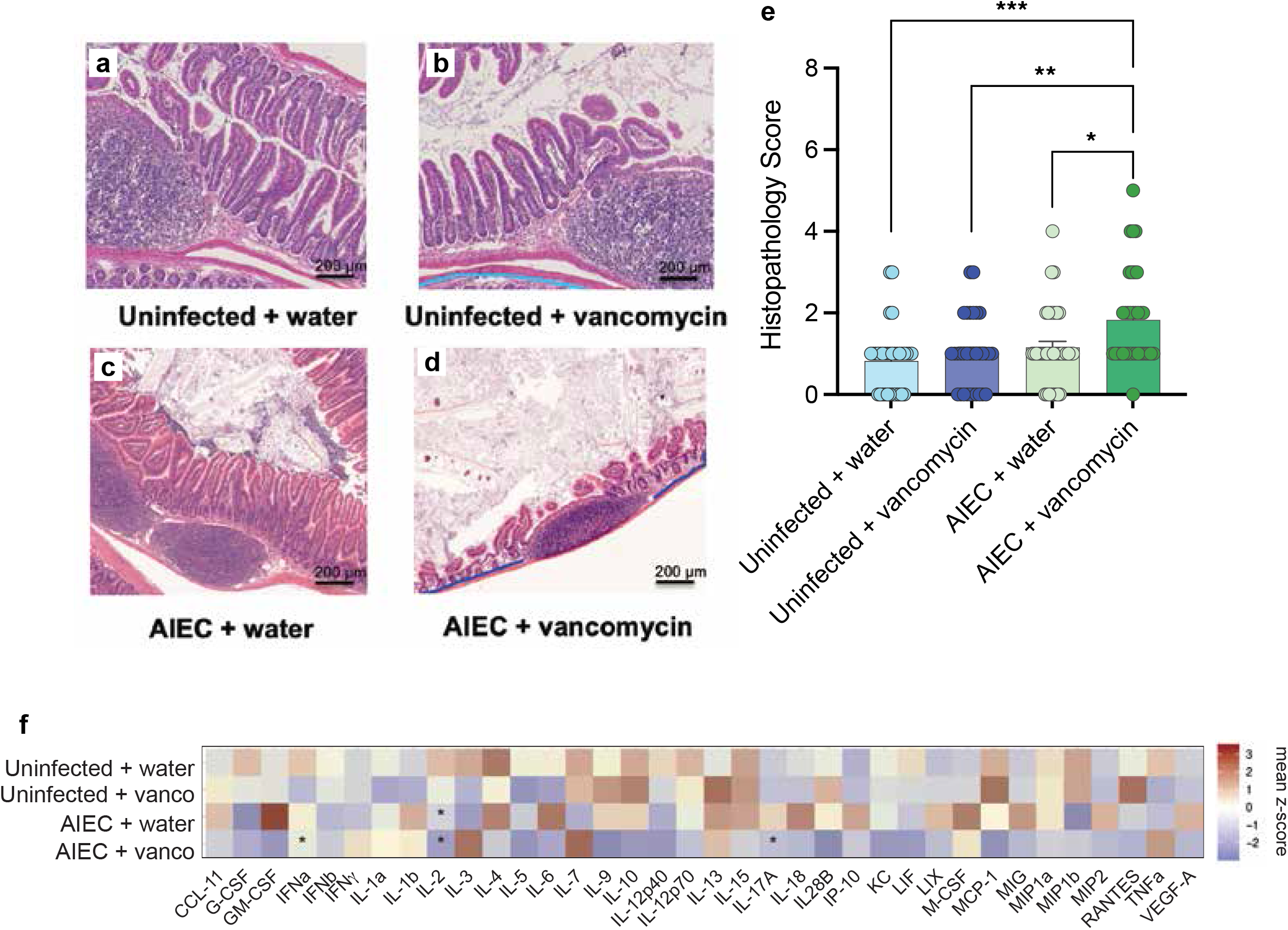
Vancomycin-associated AIEC expansion promotes focal inflammation in Peyer’s patches. Representative H&E-stained images of ileal Peyer’s patches from **(a)** uninfected water-treated mice, **(b)** uninfected vancomycin-treated mice, **(c)** AIEC-colonized water-treated mice, and **(d)** AIEC-colonized vancomycin-treated mice. Scale bars = 200 μm. **(e)** Histopathology scores of Peyer’s patch-associated regions from each treatment group (Kruskal–Wallis test with Dunn’s multiple-comparison correction). Each point represents an individual mouse. Bars represent the mean ± SEM. *p < 0.05, **p < 0.01, ***p< 0.001. **(f)** Heatmap showing mean z-score–normalized cytokine and chemokine expression profiles within Peyer’s patches across treatment groups, as determined by multiplex cytokine analysis. Asterisks indicate analytes with statistically significant differences relative to uninfected water-treated controls (Kruskal–Wallis test with Dunn’s multiple-comparison correction).

### AIEC virulence factors differentially contribute to colonization and inflammation in Peyer’s patches

We next sought to identify mechanisms that enable AIEC to expand and promote focal inflammation within ileal Peyer’s patches. Given that inflammation in this niche was specifically associated with AIEC expansion, we hypothesized that AIEC virulence factors contribute to this phenotype. To test this, mice were colonized with Δ*fliC*, Δ*lpf*, or Δ*PI-6* mutant AIEC strains and treated with either water or vancomycin. These mutants were selected based on their roles in motility (flagellin), adhesion (long polar fimbriae), and antimicrobial peptide resistance (31), respectively. Following vancomycin treatment, all three mutant strains expanded to levels comparable to wild-type AIEC in the lumen and bulk ileal tissue (**Fig. 5 a,b**). In Peyer’s patches, however, distinct differences in colonization were observed (**Fig. 5c**). The Δ*lpf* mutant colonized Peyer’s patches efficiently, while the Δ*PI-6* mutant showed reduced expansion relative to wild type, although this did not reach statistical significance. In contrast, the Δ*fliC* mutant was undetectable in Peyer’s patches, indicating that motility is required for colonization of this niche. Although long polar fimbriae and antimicrobial peptide resistance genes had limited effects on AIEC colonization of Peyer’s patches, we assessed whether these factors contribute to the focal inflammation observed in AIEC-expanded mice. Histopathological analysis revealed that all three mutant strains were associated with significantly reduced tissue pathology surrounding Peyer’s patches compared to wild-type AIEC (**Fig. 5d-h**). Cytokine and chemokine profiling showed similar trends, with Δ*lpf*- and Δ*PI-6*–colonized Peyer’s patches displaying altered inflammatory profiles relative to wild type (**Fig. 5i**). Specifically, these mutants were associated with reduced levels of IFN-α and the chemokine KC, and the Δ*PI-6* mutant also showed a distinct reduction in IL-2. In contrast, cytokine and chemokine profiles in Peyer’s patches colonized with the Δ*fliC* mutant were not significantly different from wild type.

**Figure 5.**
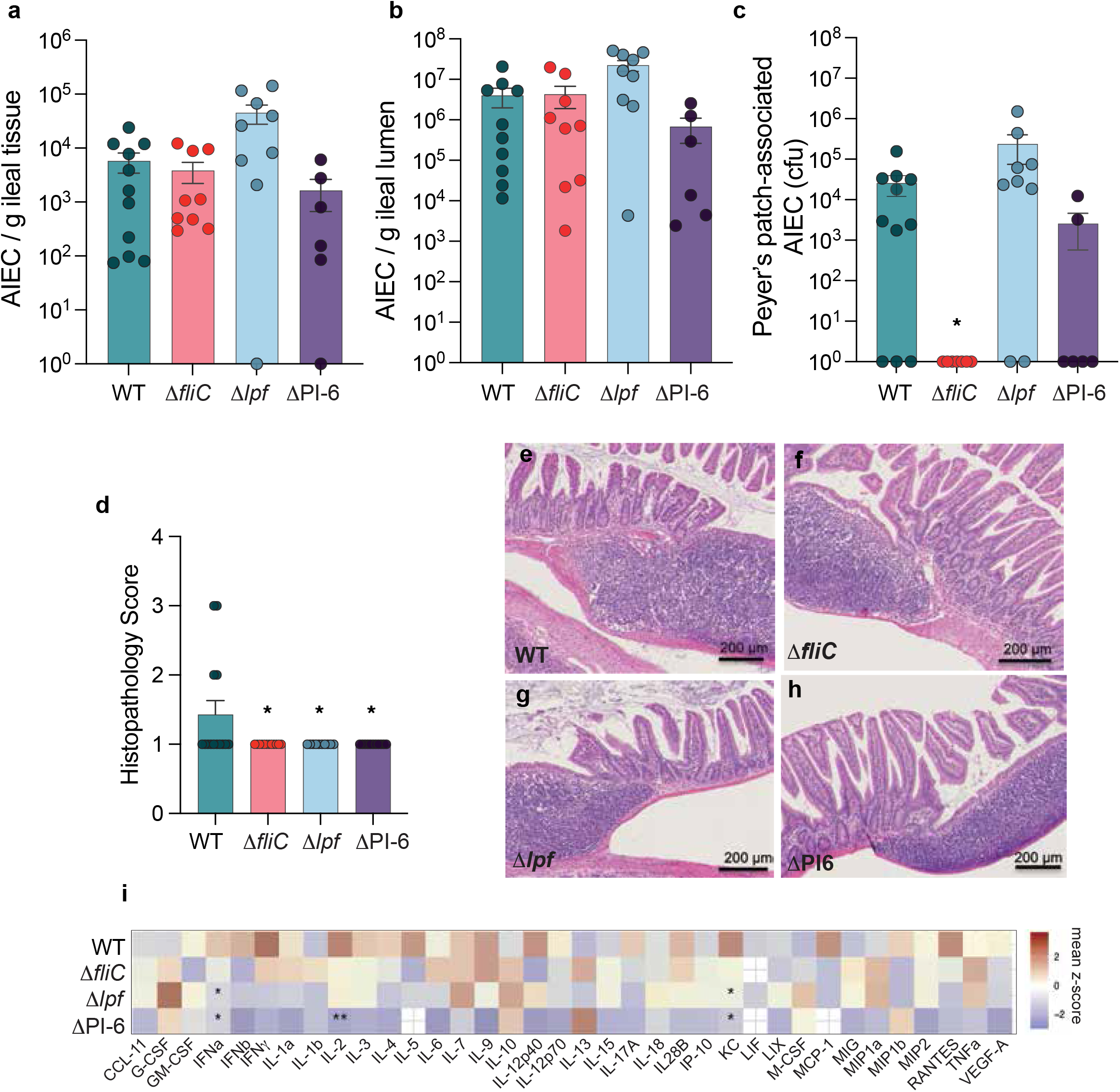
AIEC virulence factors differentially contribute to colonization and inflammation in Peyer’s patches. Mice were colonized with wild-type (WT) AIEC or the indicated mutant strains (Δ*fliC*, Δ*lpf*, or ΔPI-6) and treated with vancomycin (50 mg/kg) daily for five days. At the experimental endpoint, ileal tissue, ileal luminal contents, and Peyer’s patches were harvested to quantify bacterial burden. AIEC burden in **(a)** ileal tissue, **(b)** ileal lumen, and **(c)** Peyer’s patches following colonization with each strain (Kruskal–Wallis test with Dunn’s multiple-comparison correction). **(d)** Histopathology scores of Peyer’s patch-associated regions from mice colonized with each AIEC strain (Kruskal–Wallis test with Dunn’s multiple-comparison correction, p = 0.0180). Each point represents an individual mouse. Bars represent the mean ± SEM. Representative H&E-stained images of Peyer’s patch-associated regions following colonization with **(e)** WT AIEC, **(f)** Δ*fliC*, **(g)** Δ*lpf*, or (h) ΔPI-6 AIEC strains. Scale bars = 200 μm. **(i)** Heatmap showing mean z-score–normalized cytokine and chemokine expression profiles within Peyer’s patches colonized with WT or mutant AIEC strains, as determined by multiplex cytokine analysis. Asterisks indicate analytes with statistically significant differences relative to WT AIEC (Kruskal–Wallis test with Dunn’s multiple-comparison correction). *p < 0.05, **p < 0.01.

Taken together, these findings indicate that AIEC virulence factors differentially contribute to colonization and inflammation within Peyer’s patches, with flagellin required for niche colonization and long polar fimbriae and antimicrobial peptide resistance determinants contributing to the inflammatory response.

## DISCUSSION

Antibiotic treatment is known to promote dysbiosis and an outgrowth of CD-associated Enterobacteriaceae, including AIEC. However, it has remained unclear whether antibiotic exposure creates mucosal niches that support AIEC expansion, particularly at sites of disease activity such as Peyer’s patches. In this study, we found that vancomycin treatment promotes AIEC expansion in ileal Peyer’s patches, which is associated with focal inflammation at this compartment. Peyer’s patches are a well-established niche for enteric pathogens, as *Salmonella* Typhimurium and *Shigella flexneri* are known to infiltrate and persist in this lymphoid tissue (32–34). AIEC have also been shown to colonize Peyer’s patches using long polar fimbriae following infection of exteriorized murine ileal loops (16), however, the dynamics of Peyer’s patch colonization by resident AIEC following antibiotic treatment have not been described.

Colonization resistance is a key barrier that prevents Enterobacteriaceae expansion in luminal and mucosal niches of the GI tract (35). By perturbing the microbiome, antibiotic exposure can reduce colonization resistance to enable the expansion of these microbes (35). Consistent with this paradigm, our microbiome analysis revealed that vancomycin treatment markedly restructures bacterial composition within Peyer’s patches, with one of the most notable changes being the contraction of segmented filamentous bacteria (SFB). SFB are known to exert colonization resistance against invading pathogens in mucosal niches, as they preferentially localize to the ileal mucosa and are often found near Peyer’s patches (36) . SFB also support colonization resistance by stimulating IgA production and reinforcing mucosal immunity (37).

In parallel with AIEC expansion in Peyer’s patches following antibiotic treatment, we observed histological evidence of inflammation around this niche. This effect appeared to be specific to AIEC, as expansion of resident *E. coli* in antibiotic-treated naïve mice did not significantly affect histopathology scores, nor did infection with AIEC lacking key virulence factors that distinguish them from commensal *E. coli*. The focal inflammation observed in AIEC-expanded mice coincided with a depletion of IL-17A. As a Th17 cytokine, IL-17A maintains intestinal barrier integrity and induces antimicrobial peptide production in the lamina propria. Reduced IL-17A would therefore be expected to impair mucosal defenses, consistent with the elevated AIEC burden and tissue pathology we observed in this niche after vancomycin treatment. This finding aligns with a previous report of impaired Th17 activity following vancomycin exposure (38), an effect thought to result from depletion of SFB (37). Supporting this model, other studies of AIEC expansion have also reported reductions in Th17-associated cytokines, including IL-22 (39). Together, these data suggest that disruption of Th17-mediated immunity following antibiotic treatment may limit host control of AIEC in ileal Peyer’s patches.

Interestingly, AIEC-expanded mice also displayed a reduction in IFN-α and IL-2 within Peyer’s patches. These cytokines have not been widely implicated in Peyer’s patch dysfunction or AIEC-associated pathology, suggesting they may be a unique signature of AIEC expansion within this niche. Although IFN-α and IL-2 are predominantly pro-inflammatory cytokines, both have context-dependent roles in maintaining immune homeostasis. Type I contribute to epithelial barrier integrity and reduce susceptibility to DSS colitis in mouse models (40), while IL-2 promotes regulatory T cell responses and a tolerogenic microbiome (41). The reduction of these cytokines in therefore consistent with the increased inflammation observed in AIEC-expanded mice. However, given the pleiotropic nature of these pathways, further work will be required to define their specific roles in AIEC-driven pathology with Peyer’s patches.

We found that virulence factors contribute to AIEC expansion in Peyer’s patches, particularly the flagellin protein encoded by *fliC*. Given that motility is generally required for mucosal colonization, we expected reduced AIEC burden in both bulk tissue and in Peyer’s patches. However, deletion of *fliC* attenuated AIEC expansion in Peyer’s patches but had no effect on expansion in bulk tissue. This suggests that motility is specifically required for colonization of this niche rather than general tissue access. One possibility is that loss of flagellin reduces antigen sampling via TLR5, thereby limiting AIEC uptake into Peyer’s patches. Alternatively, deletion of *fliC* may alter the expression of other virulence factors, as flagella gene expression is often co-regulated with adhesins (42, 43). In AIEC strain LF82, deletion of the *fliA* regulator gene reduces type 1 pilus expression and impairs epithelial adhesion (44). It is therefore possible that deletion of *fliC* indirectly reduces expression of other virulence factors required for Peyer’s patch colonization.

Unexpectedly, long polar fimbriae were dispensable for AIEC expansion within Peyer’s patches, as the Δ*lpf* mutant colonized this niche to a similar extent as wild type. This contrasts with previous work showing that long polar fimbriae facilitate M cell translocation (16). Those studies used an acute ileal loop model, whereas our study examines population dynamics during established *in vivo* colonization. It is possible that other virulence factors, such as FimH, can mediate AIEC colonization of Peyer’s patches independently of long polar fimbriae. Like long polar fimbriae, FimH can also interact with M cell receptors (45, 46). These findings suggest that AIEC may use redundant or context-dependent mechanisms to access Peyer’s patches.

We found that AIEC required an intact virulence repertoire to elicit focal inflammation in Peyer’s patches, as deletion of *fliC, lpf*, or PI-6 significantly attenuated tissue pathology in this niche. This effect cannot be explained solely by differences in bacterial burden, as the Δ*lpf* mutant colonized Peyer’s patches, but resulted in less pathology. Instead, these findings suggest that specific virulence factors modulate the host immune response. Consistent with this interpretation, Peyer’s patches colonized with the Δ*lpf* and ΔPI-6 mutants displayed reduced levels of IFN-α and KC, which are associated with Th1 activation and neutrophil recruitment (47, 48). These data indicate that the determinants of colonization and inflammation within Peyer’s patches are at least partially separable.

The inflammatory profile associated with AIEC expansion in Peyer’s patches supports a role for this niche in CD pathogenesis. Peyer’s patches have long been proposed as sites of early disease activity, as mucosal lesions frequently localize near these structures (49, 50). Our findings suggest that antibiotic-induced disruption of this compartment may facilitate localized expansion of pathobionts, contributing to focal inflammation at disease-relevant sites. Notably, while resident *E. coli* was detectable in Peyer’s patches following antibiotic treatment, they did not induce pathology, highlighting the importance of pathogenic traits in driving inflammation. It is possible that resident strains may adapt within this niche over time to acquire AIEC-like properties, as observed in models of pathobiont evolution (30).

Overall, our study identifies Peyer’s patches as a site where antibiotic-induced dysbiosis enables AIEC expansion, colonization, and localized inflammatory responses. These findings highlight the importance of considering tissue-specific niches in host-microbe interactions that underly disease initiation and progression, and suggest that Peyer’s patches may represent a critical site for pathobiont persistence in CD.

## ACKNOWLEDGEMENTS

We thank the staff at the Centre for Advanced Light Microscopy for their help with obtaining high-quality images of AIEC within Peyer’s patches. K.R.I. was supported by an Ontario Graduate Scholarship and a Canadian Institutes of Health Research Canada Graduate Scholarship. This work was funded by grant PJT-470165 from the Canadian Institutes of Health Research and Crohn’s and Colitis Canada Grant-In-Aid of Research program.

